# Semaglutide Increases Striatal Leptin Sensitivity, Impacting Dopaminergic Response to Cocaine

**DOI:** 10.64898/2026.10.16.759697

**Authors:** James Muldowney, Madison Umlauf, Miriam E. Bocarsly

## Abstract

There is accumulating evidence that the GLP-1 analog, semaglutide, might effectively curb intake of cocaine via brain dopaminergic functioning, despite limited brain penetrance. However, there is currently no established mechanism elucidating how chronic semaglutide treatment regulates the dopaminergic response to cocaine. Cocaine is a highly addictive substance that produces rewarding effects primarily through antagonism of the dopamine transporter (DAT). Therapeutics for cocaine use disorder (CUD) have long targeted DAT in the hope of reducing cocaine intake. Metabolic hormones such as leptin and insulin have been shown to influence DAT, and for that reason have been explored for the regulation of cocaine reward and behavior. Interestingly, while semaglutide does not penetrate the brain, it does peripherally modulate metabolic hormones that enter the brain. Through fast-scan cyclic voltammetry and behavioral experimentation, we examined the effects of chronic semaglutide treatment on dopamine signaling, cocaine-induced dopamine release, and locomotor sensitivity. We determined that chronic semaglutide inhibits dopaminergic release induced by cocaine. Surprisingly, semaglutide’s effects on cocaine reward were blunted in mice fasted overnight, suggesting a mechanistic dependence on metabolic hormones. Semaglutide administration generated significant reductions in fasting blood leptin, as well as enhanced striatal leptin sensitivity compared to vehicle controls. Preceding cocaine exposure, application of leptin to fasted mice restored semaglutide’s regulation of cocaine. Taken together, we propose a novel mechanism underlying semaglutide’s effects on cocaine-induced dopamine, emphasizing the potential of central leptin signaling as a therapeutic target for CUD.

## Introduction

Nearly 50 million Americans suffer from a substance use disorder, with 5 million reported users of cocaine and 1.3 million patients of cocaine use disorder [1]. Interestingly, there is accumulating evidence that GLP-1 analogs, initially marketed for diabetes and weight loss, might also work to curb intake of substances, including cocaine [23]. However, much of this research was focused on earlier-generation GLP-1 receptor agonists, and the mechanisms underlying these effects in the highly efficient next-generation GLP-1 analogs, like semaglutide, remain unclear.

Research on the blood-brain penetrant glucagon-like peptide 1 receptor agonist exendin-4 and exenatide exhibits their ability to alter the reward system [24–26]. Studies have shown these analogs effectively reduce self-administration of cocaine, cocaine-induced locomotion, cocaine conditioned place preference, and cocaine-induced evoked dopamine in rodents [26–28]. Despite these findings, exenatide did not significantly decrease cocaine self-administration in CUD patients [29].

The development of a more potent GLP-1RA, semaglutide, has reinitiated the examination of GLP-1 analogs for CUD treatment. Chronic semaglutide treatment causes significant metabolic changes, such as a reduction in fasting blood leptin concentration in obese and lean models [30]. Though semaglutide demonstrates limited CNS access compared to previous analogs [31–34], with no evidence of reaching deeper midbrain regions such as the striatum, new findings reveal modified dopamine dynamics and reward behavior.

Despite its inability to penetrate the blood-brain barrier, acute semaglutide treatment increases dopamine release in the striatum during food reward while reducing eating behavior [35]. Further, chronic and acute semaglutide administration reduces alcohol intake and relapse behavior in male and female rats through regulating alcohol-induced dopamine release [36]. A case study on an obese patient receiving semaglutide treatment for weight loss detailed a substantial decline in cocaine craving [37]. Recently, acute semaglutide administration was shown to decrease cocaine self-administration and cocaine-induced dopamine signaling through microdialysis and fiber photometry [38]. Currently, there is no established mechanism elucidating how chronic semaglutide treatment regulates the dopaminergic response to cocaine, especially with limited brain penetrance.

Cocaine’s addictive properties can be attributed to its affinity for the dopamine transporter (DAT), blocking synaptic reuptake of dopamine [2–4]. Modulators of DAT functionality and models of altered striatal DAT expression emphasize the potential of DAT as a therapeutic target for CUD [5–10]. With the discovery of metabolic hormones’ influence on dopamine release and reuptake [11–13], the signaling pathways of insulin, leptin, and GLP-1 have been explored for the regulation of cocaine reward and behavior [13–19]. Further, semaglutide is known to affect peripheral hormone levels of insulin and leptin, which do effectively cross the blood-brain barrier. This led us to hypothesize that semaglutide could exert its overall effects on striatal dopamine function through its modulation of peripheral hormones such as insulin and leptin. In this study, we examined the effects of chronic semaglutide treatment on dopamine signaling, as well as semaglutide’s control over cocaine-induced dopamine release and locomotive behavior. We further investigated the mechanism of cocaine response regulation through evaluating the necessity of leptin signaling, emphasizing the potential of central leptin signaling as a therapeutic target for CUD.

## Methods

### Mice

Adult C57BL/6J male mice (JAX, 12-20 weeks) were group housed in ventilated cages (Nexgen System, Allentown Inc., PA, USA) with controlled temperature, humidity, and light (21 ± 2 °C, humidity 60 ± 10 %, 12:12 h light/dark cycle) with free access to food and water, unless specified. Mice were randomly assigned to one of two treatment groups: semaglutide (Novo Nordisk; 40 μg/kg s.c.) or vehicle (0.4% DMSO in saline (0.9% NaCl), 10 mL/kg, s.c.). Injections were given every three days for five weeks.

Fast Scan Cyclic Voltammetry (FSCV)

#### Brain slice preparation

Mice were deeply anesthetized with isoflurane, decapitated, and brains were rapidly harvested and placed in warm cutting solution (in mM): 225 sucrose, 13.9 NaCl, 26.2 NaHCO_3_, 1 NaH_2_PO_4_, 1.25 glucose, 2.5 KCl, 0.1 CaCl_2_, 4.9 MgCl_2_, and 3 kynurenic acid. Coronal brain slices were made using a vibratome (230 μm) in warm cutting solution. Brain sections containing dorsomedial striatum or the nucleus accumbens were allowed to recover in oxygenated ACSF containing (in mM): 124 NaCl, 2.5 KCl, 2.5 CaCl_2_, 1.3 MgCl_2_, 26.2 NaHCO_3_, 1 NaH_2_PO_4_, and 20 glucose (∼310-315 mOsm) at 33⁰C for 30-60 min and maintained at room temperature until recording.

#### Recordings

Fast-scan cyclic voltammetry was performed in the dorsomedial striatum of semaglutide and vehicle-treated mice to determine levels of evoked dopamine. Carbon fibers (7μm diameter) were inserted into glass pipettes to create carbon fiber electrodes (CFEs), which were hand-cut to expose ∼150μm of the fiber past the pipette tip. Before use, CFEs were backfilled with 3M KCl internal solution. CFEs were conditioned before recordings with a triangular voltage ramp (-0.4 to 1.2 and back to -0.4V versus Ag/AgCl reference at 400V/s) delivered every 15ms. During recordings, CFEs were held at -0.4V versus Ag/AgCl while the same triangular voltage ramp was delivered every 100ms. Dopamine transients were evoked by a single pulse (0.2ms) of electrical stimulation administered every 2 minutes by a glass micropipette filled with ACSF that was placed near the CFE tip. To generate input-output curves, the intensity of electrical stimulations was varied (10, 20, 40, 60, 80, 100, 150, 200, or 300µA) with each episode. Experiments were conducted in ACSF and maintained at 31-33°C. In a subset of experiments, slices were treated with semaglutide (Medex, GLP-1 receptor agonist, 100 nM), GBR12909 (Cell Signaling, DAT inhibitor, 5µM), ML375 (Millipore Sigma, muscarinic receptor 5 antagonist, 10µM), Dihydro-β-erythroidine hydrobromide (DhβE) (Tocris, nAchR inhibitor, 1µM), cocaine hydrochloride (Cayman Chemical, 1 µM and 10 µM), porcine insulin (Medex Biochemica, 100nm), or recombinant mouse leptin (R&D Systems, 100nM), all in aCSF. Data were acquired and analyzed using a Chem-Clamp amplifier (Dagan) and I/O board (National Instruments) with custom-written software VIGOR using Igor Pro (Wavemetrics) running mafPC (courtesy of M.A. Xu-Friedman). All data were analyzed with VIGOR and peak current amplitude of dopamine was converted to concentrations according to the post-experimental calibration.

### Locomotor Activity

Locomotion testing was conducted in the light cycle either in a fasted (16 hours) or fed state, as indicated in the data. Mice were transferred to the experimental room and allowed to habituate for 30 minutes, then transferred into polycarbonate chambers (20 cm H x 17 cm W x 28 cm D) equipped with infrared photobeam detectors (Columbus Instruments). Baseline locomotion was recorded for 1 hour, and 1.5 hours for leptin. On days 1-3, fed mice were then treated with saline (10 mL/kg, i.p.), and locomotor activity was collected for an additional hour. On days 4 and 5, instead of saline, mice received either the DAT inhibitor GBR12909 (i.p., 1, 10 mg/kg, each day respectively), cocaine (i.p. 10 mg/kg), or the D1-like agonist SKF-81297 (i.p., 5, 7.5 mg/kg). Mice used for fasted locomotion experiments received saline in a fed state (for habituation to injections and handling) on days 1 and 2, then saline in a fasted state on days 3 and 5. Drugs were administered to fasted mice on days 7 and 9. Data was extracted and analyzed using IGOR Pro (64-bit).

### Western blot

Mice were euthanized using isoflurane and dorsal striatum samples were taken. Samples were flash frozen and stored at −80°C. Each tissue sample was lysed in 150 μL of RIPA buffer with protease inhibitor (Thermoscientific 89901), homogenized by a hand homogenizer, and then sonicated for 15 minutes. Protein was estimated using a Pierce BCA protein assay (Thermofisher 23225) at a 1:5 dilution. 20 µg of protein from each sample was boiled for 7 minutes at 95 degrees Celsius in loading buffer. Samples were loaded onto TGX stain-free mini gels (Bio-Rad) and run at 100 V for ∼2 h. Gels were transferred to a 0.2 µm nitrocellulose membrane using the Bio-Rad Trans-Blot Turbo at 2.5 amps, 25 V for 6 min. Ponceau staining was utilized to determine equal protein loading across wells. Membranes were blocked in 1% BSA in tris-buffered saline solution with tween (TBST) for one hour at room temperature with rapid shaking.

Primary antibodies (listed in Table 1) were diluted in TBST and incubated at 4°C with gentle shaking. Following primary incubation, membranes were rinsed with TBST 3×for 10 min with shaking and then incubated with secondary HRP (Gt, Thermo Fisher, 18-4818-82, 1:5000 or Rb, Thermo Fisher, NA9340V, 1:5000) in 1% BSA in TBST for 2 hours at room temperature with gentle shaking, followed by 3×rinses with TBST for 10 min each. To develop, membranes were covered in ECL (GE Healthcare, 17030484) for 2 min and imaged using chemiluminescence on the Chemi-doc imaging system. In addition to using the stain-free feature to ensure equal loading, actin was used as a loading control. For blots in which multiple proteins were detected from a single membrane, membranes were stripped between hybridizations with a stripping buffer consisting of 20 mL 10% SDS and 12.5 mL of Tris HCl, pH 6.5, with 0.8% beta-mercaptoethanol added immediately prior to stripping. Membranes were placed in stripping buffer for 15 min shaking at room temperature. Membranes were moved to TBST and rinsed 3×for 10 min each. Membranes were re-imaged to ensure adequate stripping before re-blocking. Primary and secondary incubations were repeated as described. Band intensity was quantified using the Image Lab software (Bio-Rad).

**Table 1:**

| Protein | Antibody | Weight (kDa) | Dilution |
| --- | --- | --- | --- |
| Actin | Abcam Ab8827 | 42 | 1:1000 |
| Muscarinic Acetylcholine Receptor 5 | Abcam Ab186830 | 61 | 1:500 |
| Tyrosine-Hydroxylase | Millipore AB152 | 62 | 1:1000 |
| Phospho-TH (ser40) | Abcam ab137869 | 62 | 1:1000 |

### Metabolic assays

Mice were fasted (16 hours), and tissue was collected. Blood glucose was determined via tail blood using a handheld glucometer (Contour Next One). Trunk blood was collected in non-heparinized capillary tubes, centrifuged at room temperature and serum was collected. Serum concentrations of leptin (EZML-82K, Millipore), insulin (#90080, Crystal Chem), and adiponectin (#80569, Crystal Chem) were determined via commercially available ELISAs. Fat pads were dissected bilaterally and weighed; eWAT was slightly cranial to the epididymis.

### Statistics and analysis

Analyses were performed in Prism 10 (GraphPad). Data making three or more comparisons were analyzed using two-way ANOVA, with the addition of repeated-measures (RM) or a mixed-effects model, as appropriate (ex. voltammetry input/output curves and time courses, locomotor behavior, etc.). For RM-ANOVA, sphericity was assessed with Mauchly’s test. Significant main effects or interactions were followed up with pairwise tests corrected for multiple comparisons (Sidak or Tukey). Linear mixed effects models were utilized when multiple tissue slices from a single animal were examined. Data comparing two groups (ex. blood analysis, protein levels, etc. between treatment groups) were analyzed with an unpaired Student’s t-test. Mann-Whitney tests were run when comparing groups that failed the Shapiro-Wilk test for normality. One-sample t-test was used to determine deviations from the mean or baseline. Results were considered significant at an alpha ≤ 0.05. All data are presented as mean ± SEM, and individual animal data are shown whenever possible. Full statistics are presented in the figure legends. Sample sizes were chosen based on previous research employing similar approaches and are sufficient for detecting effects while using the minimal number of animals. FSCV experiments with drug applications were normalized to steady-state baseline values preceding drug introduction to the well. Locomotor experiments were normalized to the group average locomotion during saline control trials preceding drug injection days.

## Results

### Chronic semaglutide treatment alters behavioral and dopaminergic response to cocaine

To model clinical, chronic use of semaglutide, mice were injected with semaglutide (40 µg/kg, s.c.) every 3 days for five weeks [30,45,46]. The control group received vehicle injections on the same schedule. Following treatment, sensitivity to an acute dose of cocaine (10 mg/kg, i.p.) was tested using locomotor response (Figure 1A). While no differences were seen in locomotion in response to a novel environment (Supplementary Figure 1A) or following a saline injection (Supplementary Figure 1B), mice previously treated with semaglutide showed a blunted locomotor response to cocaine compared to vehicle-treated controls (Figure 1B). Because diminished locomotor responses to cocaine suggest alterations in striatal dopamine transmission [47], we assessed striatal dopamine release capacity. Following semaglutide or vehicle treatment, we used ex vivo fast-scan cyclic voltammetry to measure dopamine signals evoked by electrical stimulation in striatal tissue. We focused our recordings on the DMS given its distinct dopamine dynamics and established roles in cocaine seeking, value-guided action selection, and behavioral flexibility [39–44, 48–50]. We found that mice previously treated with semaglutide had a blunted dopaminergic response to cocaine (1 µM) compared to controls (Figure 1D,E). Interestingly, the decay of the dopamine signals, as determined by the decay time constant tau, was significantly faster in mice treated with semaglutide compared to controls (Figure 1F). Across all mice, we observed a correlation in evoked dopamine release and cocaine-induced inhibition of dopamine (Figure 1G), emphasizing the role of dopamine reuptake for regulating the dopaminergic response to cocaine.

**Figure 1:**
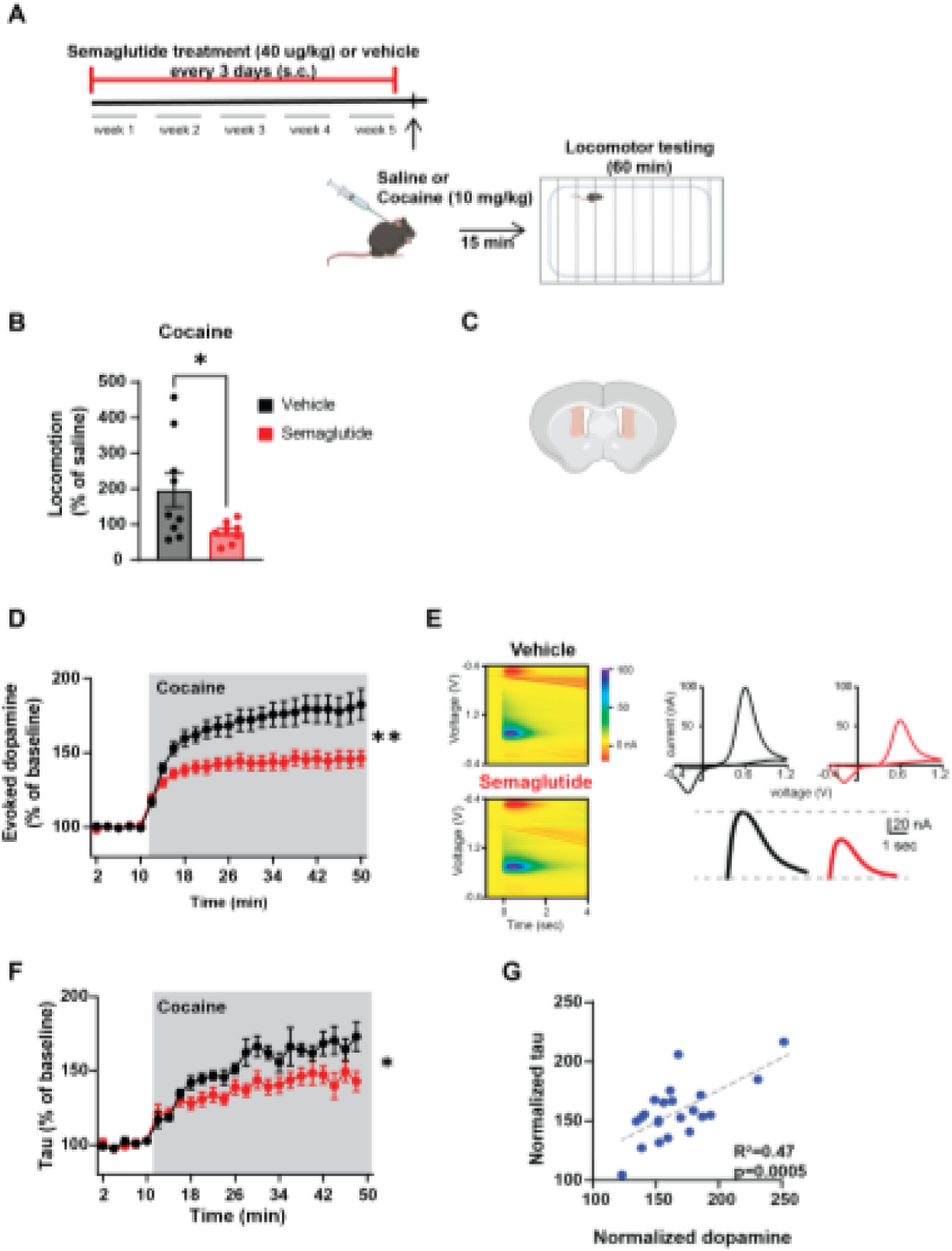
Chronic semaglutide treatment alters cocaine response and striatal dopamine dynamics in postprandial conditions. (**A**) Schematic demonstrating injection protocol and experimental design. (**B**) Average locomotor counts in cocaine-injected (10 mg/kg) mice over 60-minute trial, normalized to group-average saline control trial (t(15)=2.25, p=0.04. n = 8-9 mice/group, unpaired t-test). (**C**) Schematic of recording site: coronal mouse brain slice, with dorsomedial striatum highlighted In red (**D**) Time course of evoked dopamine in response to cocaine (1 μM) application, normalized to steady-state baseline values (interaction effect: F(24,552) = 9.92, p < 0.0001; main effect of semaglutide treatment: F(1,23) = 10.61, p = 0 004; main effect of drug application: F(1.39, 31.95) = 115.4, p < 0.0001, n = 12-13 slices/3 mice/group. RM two-way ANOVA). (**E**) **Left:** Representative 3D voltammograms showing background-subtracted current (false color scale) as a function of time (x axis) and voltage applied (y axis). **Right**: (top) representative GV plots of the electrochemical current (background-subtracted) evoked by electrical stimulation in the DMS showing the characteristic current peaks at 0.6 V and -0.2 V that match the oxidation and reduction potentials for dopamine. (bottom) Representative traces of dopamine signals from the DMS of mice previously treated with vehicle (black) or semaglutide (red). (**F**) Time course of tau values with cocaine application (1 μM), normalized to steady-state baseline values (Interaction effect: F(24,474) = 2.56, p < 0.0001; main effect of semaglutide treatment: F(1,20) = 7.05, p = 0.02; main effect of drug application: F(24.474) = 44.62. p < 0.0001, n = 10-12 slices/3 mice/group, mixed-effects analysis). (**G**) Positive correlation between normalized dopamine and normalized tau following cocaine (1 μM) application (R^2^ = 0.47, p = 0.0005, n = 22 slices/6 mice, simple linear regression). Data is presented as mean ± SEM; symbols are individual sample data. For all panels, ’p < 0.05, **p < 0.01.

As cocaine predominantly has its actions on dopamine via blocking the dopamine transporter (DAT), preventing dopamine from being cleared [51], we evaluated the effects of the DAT inhibitor, GBR-12909 (1 mg/kg, i.p.), on locomotor output following chronic semaglutide or vehicle treatment. The semaglutide-treated mice demonstrated a diminished response to DAT inhibition compared to vehicle controls (Supplementary Figure 1D,E), mimicking the response to cocaine.

Expression and phosphorylation of tyrosine hydroxylase (TH), the rate-limiting step in dopamine production, were quantified through Western blotting and showed no changes following semaglutide treatment compared to vehicle controls (Supplementary Figure 2A). Given no changes in this dopamine synthesis parameter, to better understand how semaglutide alters cocaine-mediated dopamine release, we looked into modulators of dopamine reuptake and locomotor behavior. Previous research has shown that the muscarinic acetylcholine receptor 5 (M5) present on dopamine neuronal synapses regulates dopamine release and reuptake [52, 53], possibly contributing to the dopaminergic response to cocaine [54, 55]. Semaglutide caused no significant alteration in M5 expression (Supplementary Figure 2B) nor evoked dopamine and reuptake in response to inhibition (Supplementary Figure 2C,D). Additionally, prior studies have shown how the D1 receptor can regulate the behavioral response to cocaine [56]. We used multiple doses of SKF-12987, a D1-like receptor agonist, to assess locomotive responses and determined no differences between groups (Supplementary Figure 2E). These experiments suggest that M5 and D1 signaling are not significant for semaglutide’s control of the cocaine response.

### Evoked dopamine is elevated in mice chronically treated with semaglutide

Evaluation of baseline evoked dopamine in the absence of cocaine could grant insight into how semaglutide alters cocaine-mediated dopamine release. Interestingly, we found that evoked dopamine was elevated in mice chronically treated with semaglutide compared to vehicle (Figure 2A,B). This was accompanied by a decreased decay time constant tau (Figure 2C), indicating faster clearance of the dopamine from the synapse. In a complementary study, we determined the mean amplitude of dopamine signals evoked by electrical stimulation of increasing intensities in the dorsomedial striatum of mice chronically treated with semaglutide or vehicle (Figure 2D). This experiment replicated our previous finding, with semaglutide-treated mice showing elevated levels of evoked dopamine compared to controls.

**Figure 2:**
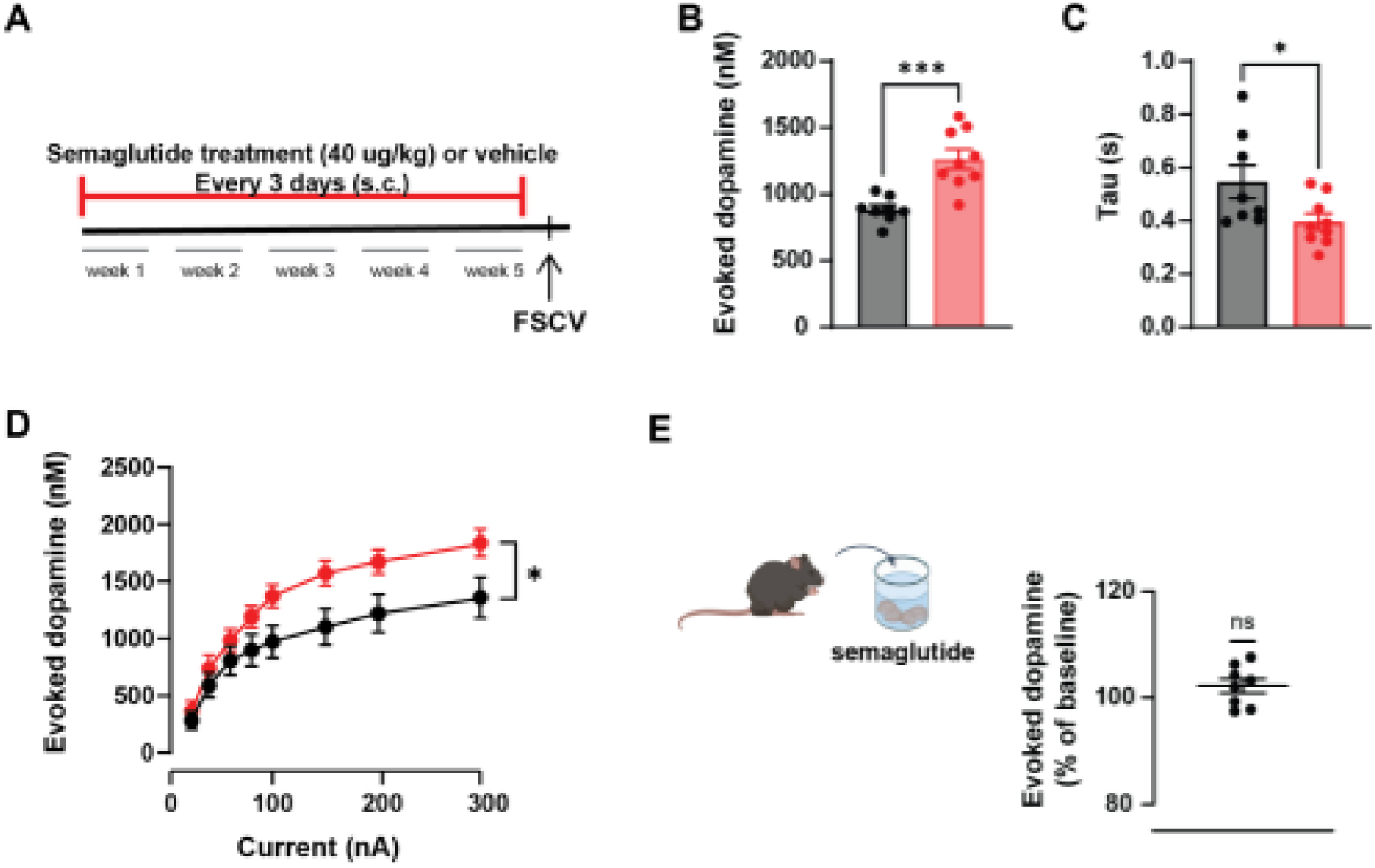
Chronic semaglutide treatment increases striatal dopamine release and reuptake. (**A**) Schematic demonstrating injection protocol and experimental design. (**B**) Average nanomolar concentration of evoked dopamine signals at 200uA stimulation in the DMS following chronic semaglutide or chronic vehicle treatment per mouse (t(15)=4.53, p = 0.0004, n = 36-41 slices/8-10 mice/group, unpaired t-test). (**C**) Corresponding values of the decay constant tau compared between semaglutide and vehicle treatment (t( 15)=2.26, p = 0.04, n = 34-37 slices/8-9 mice/group, unpaired t-test). (**D**) Evoked dopamine concentrations at varying stimulation currents in the DMS of mice chronically treated with either vehicle or semaglutide (Interaction effect: F(7,175) = 2.57, p = 0.02; main effect of semaglutide treatment: F(1,25) = 4.63, p = 0.04; main effect of current: F(1.54,38.44) = 72.65, p < 0.0001. n = 12-15 slices/3 mice/group, RM two-way ANOVA). (**E**) Evoked dopamine concentration following incubation in 100 nM semaglutide, normalized to a steady-state baseline concentration (t(7)=1.67, p = 0.14, n = 8 slices/3 mice, one sample t-test). Data is presented as mean ± SEM; symbols are individual sample data. For all panels, *p < 0.05, ***p < 0.001.

### Semaglutide does not operate directly to affect dorsomedial striatal dopamine

Again, using ex vivo FSCV, when a 100 nM concentration of semaglutide was applied directly onto brain slices containing the dorsal striatum, no changes in dopamine were detected (Figure 2E). In correspondence with previous research displaying semaglutide’s limited penetration across the blood-brain barrier, this shows that semaglutide is not working locally in the dorsal striatum to affect dopamine levels [31–34]. Because the central response does not align well with the behavioral effects, it remains unclear how this drug is influencing reward processing in deeply buried brain structures like the striatum.

### Fasting eliminates semaglutide’s regulation of cocaine response despite sustaining altered dopamine dynamics

Further experimentation revealed that the effect of semaglutide on cocaine-evoked dopamine release depended on metabolic state. As described above, mice received semaglutide or vehicle for five weeks. Prior to ex vivo FSCV, mice were fasted for 16 hours (Figure 3A). Following a 16-hour fast, cocaine-evoked dopamine release and decay time constant tau were comparable between semaglutide- and vehicle-treated mice (Figure 3B,C). Moreover, there was no significant correlation of cocaine-induced inhibition of dopamine reuptake with evoked dopamine released in each slice (Supplementary Figure 3A). Consistent with these findings, when injected with cocaine (10 mg/kg), semaglutide- and vehicle-treated mice showed similar locomotor phenotypes (Figure 3D).

**Figure 3:**
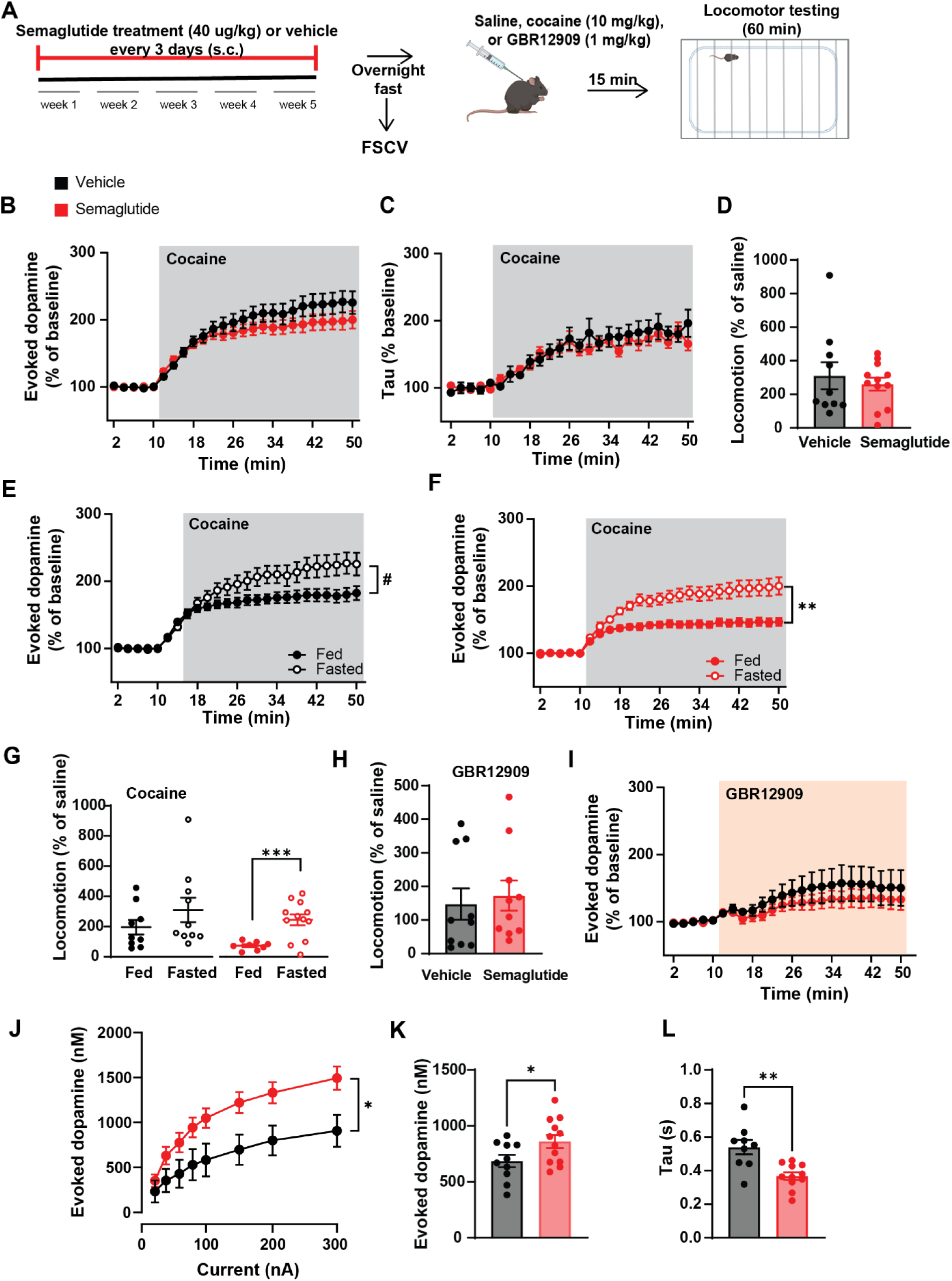
Fasting eliminates semaglutide’s regulation of cocaine response despite sustaining altered dopamine dynamics. (**A**) Schematic demonstrating injection protocol and experimental design. (**B**) Time course of evoked dopamine in response to cocaine (1 μM) application, normalized to steady-state baseline values (Interaction effect: F(24,671) = 1.93, p = 0.005; no main effect of semaglutide treatment F(1.28) = 1.22, p = 0.28; main effect of drug application: F(1.457, 40.72) = 97.94, p < 0.0001, n = 13-17 sJices/4 mice/group, mixed-effects analysis). (**C**) Time course of corresponding tau values following cocaine (1 pM) application, normalized to steady-state baseline values (no interaction effect: F(24,423) = 1.00, p = 0.47; no main effect of semaglutide treatment F(1,18) = 0.18, p = 0.31; main effect of drug application: F(4.79,84.41) = 33.66, p < 0.0001, n = 9-11 slices/4 mice/group, mixed-effects analysis). (**D**) Average locomotor counts of cocaine-injected (10 mg/kg) mice from 60-minute trial, normalized to group-average saline control tnal (vehicle Mdn = 184.3, semaglutide Mdn = 277.0, U = 59.00. z = 92.77. p = 0.97, n=10-12 mice/group, Mann-Whitney test) (**E**) Comparison of evoked dopaminergic cocaine response between fasted and fed-stale vehicle (Interaction effect: F(24,551) = 5.01, p < 0.0001; main effect of metabolic state: F(1,23) = 3.93, p = 0.06; main effect of drug application: F(1.49,34.15) = 88.52, p < 0.0001, n = 12-13 slices/3-4 mice/condition, mixed-effects analysis) and (**F**) semaglutide treated (Interaction effect: F(24,672) = 10.50. p < 0.0001: main effect of metabolic state: F(1,28) = 13.20, p = 0.001: main effect of drug application: F(1.40, 39.15) = 75.97, p < 0.0001, n = 13-17 slices/3-4 mice/condition, RM two-way ANOVA). (**G**) Comparison of cocaine-induced locomotive response between fasted and fed-state vehicle (**left**) (fasted Mdn = 184.3, fed Mdn = 126.1, U = 30.00, z = -58.14, p = 0.24, n = 9-10/condition, Mann-Whitney test) and semaglutide-treated (**right**) mice (t(18)=3.77, p = 0.001, n = 8-12/condition, unpaired t-test). (**H**) Average locomotion following GBR 12909 (1 mg/kg) from 60-minute trial, normalized to group-average saline control trial (vehicle Mdn = 96.43, semaglutide Mdn = 133.7, U = 38.00, z = 37.28, p = 0.39, n = 10/group, Mann-Whitney test). (**I**) Time course of evoked dopamine In response to 5 μM GBR application, normalized to steady-state baseline values (No interaction effect: F(23,529) = 0.52, p = 0.97; no main effect of semaglutide treatment: F(1,23) = 0.71, p = 0.41; main effect of drug application^-^ F(1.23,28.36) = 8 77, p = 0.004, n = 10-15 slices/3-4 mice/group, RM two-way ANOVA). (**J**) Evoked dopamine concentrations at varying currents in the OMS of mice previously treated with vehicle or semaglutide (interaction effect: F(7,112) = 7.40, p < 0.0001; main effect of semaglutide treatment: F(1,16) = 5.08, p = 0.04; main effect of current: F(2.14,34.23) = 113.6, p < 0.0001. n = 8-10 slices/6 mice/group. RM two-way ANOVA). (**K**) Average nanomolar concentration of evoked dopamine signals at 200 pAin the DMS per mouse (t(20) = 2.16, p = 0 04, n = 28-37 slices/10-12 mice/group, unpaired t-test). (L) Average tau per mouse compared between groups of vehicle or semaglutide treatment (t(18) = 3.69, p = 0.002, n = 21-26 süces/9-11 mice/group, unpaired t-test). Data is presented as mean ± SEM; symbols are individual sample data. For all panels, *p < 0.05,**p < 0.01,***p < 0.001.

Direct comparison of evoked dopamine amplitudes between fed and fasted control mice revealed increased cocaine-evoked dopamine release in the fasted state (Figure 3E). However, the difference in the cocaine-induced dopamine response was increased between the fasted and fed states following chronic semaglutide (Figure 3F). Consistent with these findings, fasting eliminated semaglutide’s efficacy in reducing the locomotor response to cocaine (Figure 3G).

When we replicated this study, replacing cocaine with the DAT inhibitor, GBR-12909 (1 mg/kg, i.p.), locomotor and evoked dopamine were likewise normalized across semaglutide and vehicle-treated groups following an overnight fast (Figure 3H,I).

It is important to note that while fasting abolished the dopaminergic and locomotor effects of chronic semaglutide, baseline evoked dopamine was still elevated in semaglutide-treated mice compared to vehicle-treated mice (Figure 3J-L). Together, these findings suggest that fasting may attenuate the effects of chronic semaglutide treatment on cocaine-evoked dopamine signaling and prompted us to investigate the metabolic factors that differ between fed and fasted states.

### Fasting blood leptin levels are decreased in semaglutide-treated mice

Although the metabolic effects of semaglutide treatment are well characterized in obese mice, its effects in lean mice remain less well understood. One study reported that chronic semaglutide administration alters circulating leptin levels and fat accumulation in lean mice [30]. Here, we found that chronic semaglutide treatment had no effect on body weight in lean mice (Figure 4A). Likewise, fasting blood glucose and insulin levels were not statistically different between treatment groups (Figure 4B,C). However, we did find semaglutide-induced reductions in fasting leptin concentrations (Figure 3D), similar to the previous study. As leptin is proportionally secreted to the mass of adipose tissue and predominantly regulated by visceral fat in male mice [57], epididymal white adipose tissue (eWAT) was extracted from semaglutide and vehicle-injected mice and subsequently measured to determine the ratio of fat weight to total body weight. This resulted in a decrease in the ratio in semaglutide-treated mice compared to vehicle controls (Figure 3E). We also looked at fasting blood adiponectin levels. Like leptin, adiponectin is an adipokine secreted by adipose tissue; however, circulating adiponectin levels generally vary inversely with leptin levels [58]. In the current study, while adiponectin was trending upwards following semaglutide treatment, it was not statistically significant (Figure 4F).

**Figure 4:**
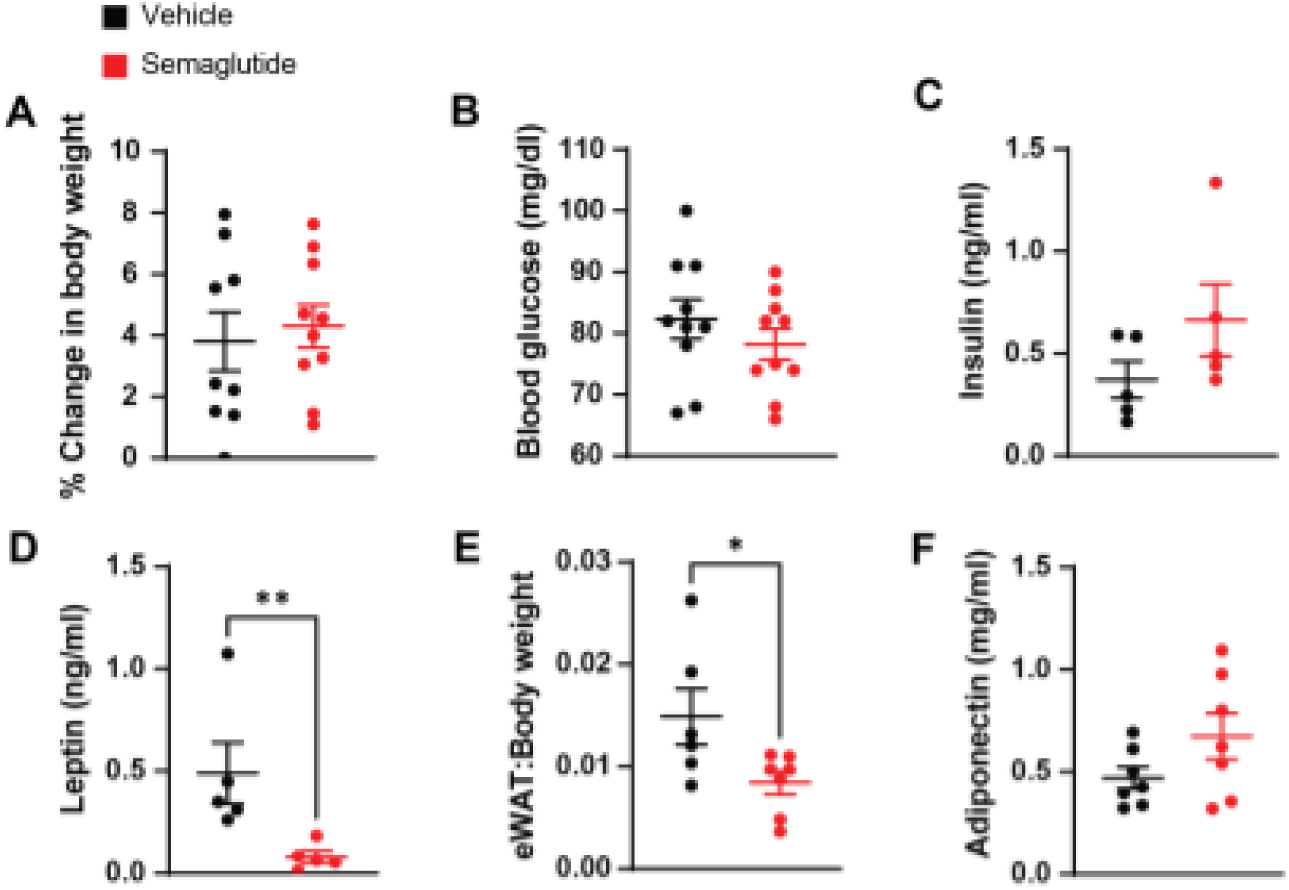
Semaglutide alters metabolic markers in lean mice without weight change. (**A**) Percent change in body weight from pre-injection period to end of 5-week treatment of vehicle or semaglutide (t(17) = 0.43. p = 0.67, n = 9-10 mice/group, unpaired t-test). (**B**) Fasting blood glucose measurements taken via tail blood at the end of the treatment period (t(18) = 1.00, p = 0.33, n = 10 mice/group, unpaired t-test). (**C**) Measurements of fasting insulin (t(8) = 1.47, p = 0.18, n = 5/group, unpaired t-test). (**D**) Measurements of fasting leptin (vehicle Mdn = 0.35, semaglutide Mdn = 0.06, U = 0.00, z = -0.29. p = 0.008, n = 5 mice /group, Mann-Whitney). (**E**) Ratio of epididymal fat pad weight to total body weight (t(11) = 2.31, p = 0 04, n = 6-7 mice/group, unpaired t-test). (**F**) Measurements of fasting adiponectin concentrations in blood serum (t(12) = 1.63, p = 0.13, n = 7 mice/group, unpaired t-test). Data is presented as mean ± SEM; symbols are individual sample data. For all panels. *p < 0.05,**p<001.

### Semaglutide induces increased striatal leptin sensitivity for the regulation of cocaine responsivity

While the reduction in peripheral leptin is not surprising because it is consistent with previous findings following semaglutide treatment [30], it identified a potential mechanistic link between peripheral metabolic changes and striatal dopamine signaling, as leptin crosses the blood–brain barrier and modulates dopamine function in the striatum [12, 53]. This led to investigating leptin’s potential significance in cocaine regulation following semaglutide treatment. Again, mice were chronically treated with semaglutide or vehicle, fasted overnight to normalize and minimize leptin levels across mice, and locomotor activity was recorded following an acute injection of leptin (Figure 5A). The literature suggests that central leptin activation following an i.p. injection occurs approximately 30 minutes after peripheral injection [59] and lasts for as long as 3 hours [12], leading us to use the last 60 minutes of the 90-minute trials for analysis. Leptin injections resulted in a significantly increased locomotor response from semaglutide-treated mice compared to vehicle controls (Figure 5B). Further investigation of the underlying dopaminergic state was conducted using FSCV. Following the application of a 100 nM leptin solution, the evoked dopamine response was greater in semaglutide-treated mice compared to vehicle controls (Figure 5C).

**Figure 5:**
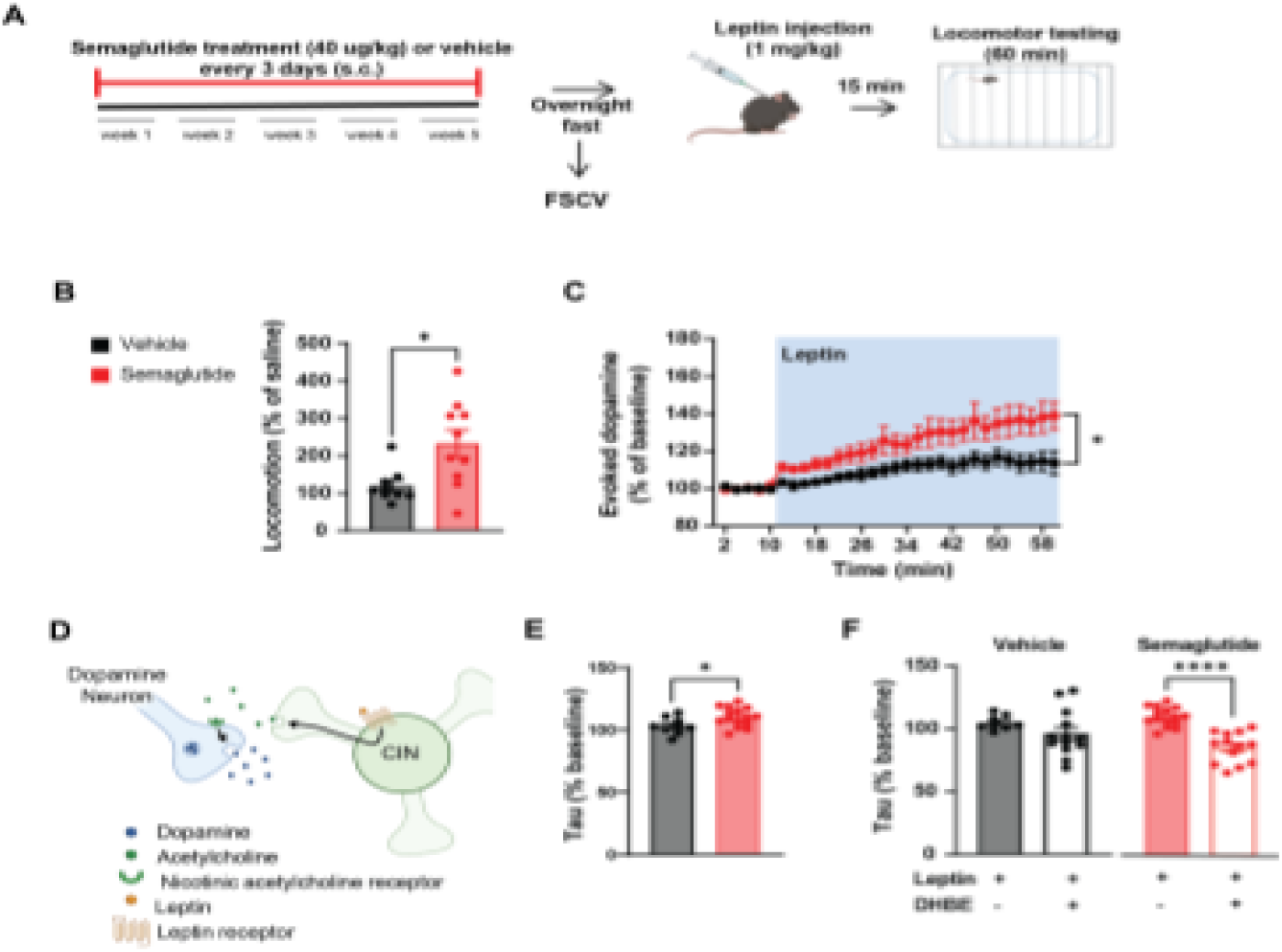
Semaglutide induces increased striatal leptin sensitivity in the dorsal striatum. (**A**) Schematic demonstrating injection protocol and experimental design. (**B**) Average locomotion of leptin-injeefed (1 mgi’kg) mice from the last 60 minutes of a 90-minute trial, normalized to group-average saline control tnal (vehicle Mdn = 107.6. semaglutide Mdn = 226.5, U = 15 00, z = 118.9. p = 0 01, n = 9-10 mice/ group. Mann-Whitney). (**C**) Normalized evoked dopamine time course in response to 100 nM leptin application (Interaction effect: F(29.654) = 3.04. p < 0.0001: main effect of semaglutide treatment: F( 1, 23) = 4.94, p = 0.04; main effect of drug application: F(1.29. 29.16) = 19.34, n = 11-14 slices/3-4 mice/group, mixed-effects analysis). (**D**) Schematic of striatal leptin control over acetylcholine-induced dopamine release tf-rough cholinergic interneuron (CIN) activation. (**E**) Normalized tau in response to 100 nM leptin application (t(22) = 2.30, p = 0.03, n = 9-15 slices/3-4 miceigroup, unpaired t-test). (**F**) Normalized tau In response to leptin compared to leptin and DhβE (1μM) in vehicle-treated mice (t(20) = 1.17, p = 0.26, n = 9-13 slices/3-4 mieoicondition, unpaired t-test) and semaglutide-treated mice (t(26) = 6.51, p < 0.0001, n = 13-15 slices/4 mice/condition). Data is presented as mean ± SEM; symbols are individual sample data. For all panels, *p <0.05, ****p< 0.0001.

Importantly, the current findings are specific to leptin and not all peripheral hormones in general. Previous research has shown that insulin regulates dopamine release in the striatum in the same fashion as leptin [60], and it has also been investigated for regulating addiction [14]. However, when insulin was applied to brain slices from mice chronically treated with semaglutide or vehicle, no changes in the dopaminergic response were observed (Supplementary Figure 4A).

### Leptin elevates dopamine reuptake in semaglutide-treated mice

Leptin influences striatal dopamine release through two distinct mechanisms: activation of cholinergic interneurons (CIN) to increase local dopaminergic release (as depicted in Figure 5D) and actions directly on the dopaminergic neurons (not shown). CIN depolarization from leptin generates acetylcholine release, subsequently binding to nicotinic receptors on dopamine neurons to increase dopaminergic firing [59]. In parallel, leptin directly influences dopaminergic neurons by elevating DAT activity [12]. With leptin application in our FSCV experiments, along with elevations in evoked dopamine concentration, the semaglutide-treated mice exhibited a significant increase in the decay constant tau (Figure 5E). This can be attributed to the expected rise in acetylcholine from leptin application, which increases dopamine release, particularly in semaglutide-treated mice. To assess leptin-induced dopamine reuptake alterations, we sought to uncouple the local CIN-regulated dopamine release and the effects of leptin on DAT functionality. To do this, we applied the nicotinic receptor antagonist, DhβE, to block the CIN contributions before the addition of leptin during FSCV. Tau values were assessed as a readout of dopamine clearance, showing that in the presence of DhβE and leptin, semaglutide-treated mice showed a significant reduction in tau, whereas vehicle-treated controls did not (Figure 5F). This demonstrates that when CIN-mediated release of dopamine is taken offline, we can clearly see semaglutide’s effects on leptin’s regulation of dopamine reuptake, which we hypothesize reflects a semaglutide-induced increase in DAT activity.

### Semaglutide reduces cocaine responsivity through enhanced dopamine reuptake mediated by leptin

We next aimed to determine the relationship between leptin and cocaine in semaglutide-treated mice. Mice that had been chronically administered either semaglutide or vehicle were fasted overnight, then given a peripheral injection of leptin, followed by cocaine, in order to assess leptin’s inhibition of cocaine-induced locomotion (Figure 6A). Locomotor activity following cocaine was normalized to locomotor response following leptin. Semaglutide-treated mice demonstrated a stark reduction in locomotion compared to controls (Figure 6B), which resembled the locomotor response to cocaine observed in a fed-state (Figure 1C). Again, we looked at evoked dopamine in the dorsal striatum of fasted mice chronically treated with either semaglutide or vehicle, this time applying leptin to the bath preceding cocaine. We found that mice previously treated with semaglutide show less evoked dopamine and increased dopamine reuptake compared to vehicle-treated mice (Figure 6C,D).

**Figure 6:**
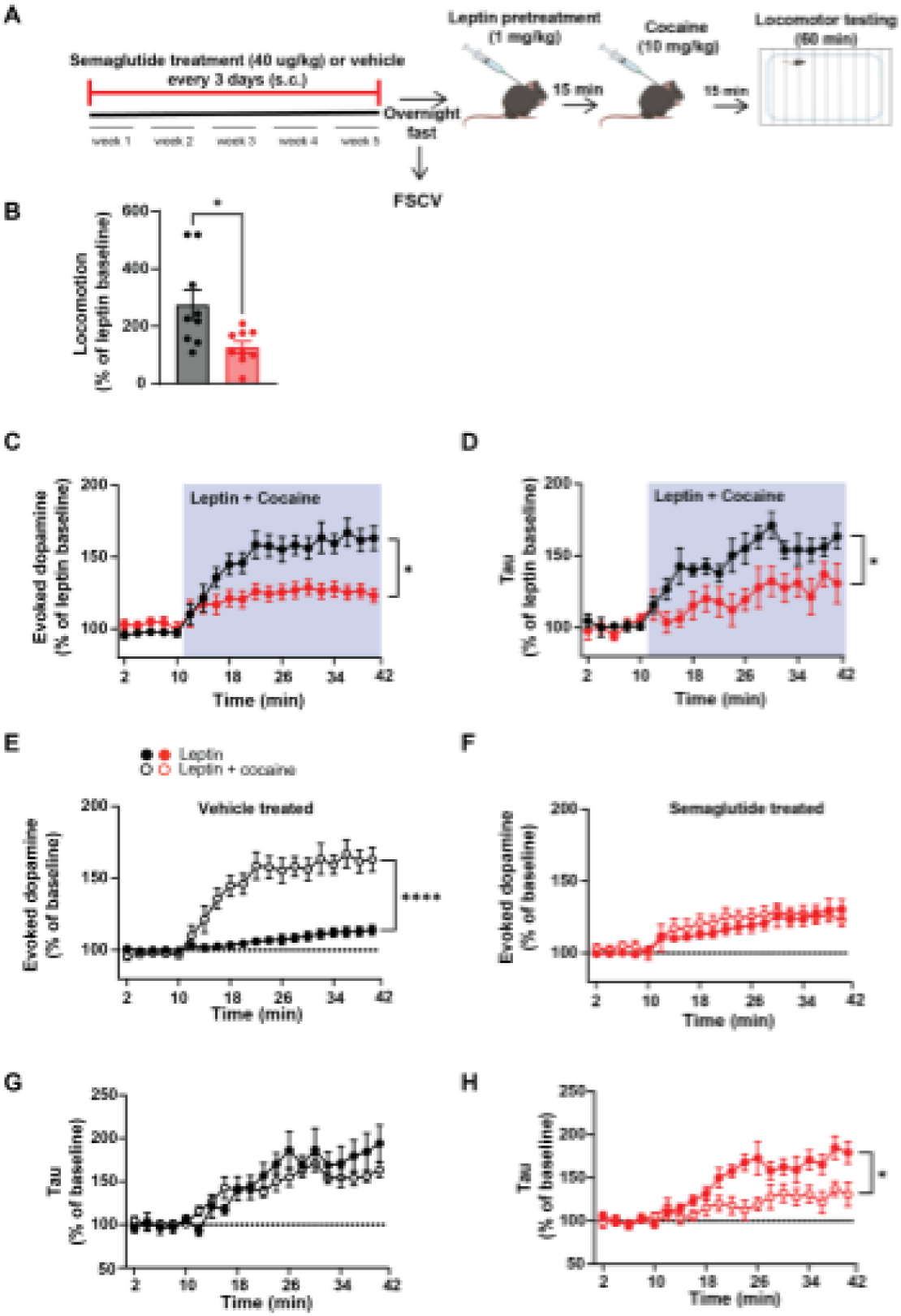
Leptin restores semaglutide’s efficacy in reducing the response to cocaine in fasted mice. (**A**) Schematic demonstrating injection protocol and experimental design. (**B**) Average locomotion of cocaine-injected (10 mg/kg) mice following leptin (1 mg/kg) pretreatment, normalized to group leptin response (t( 16) = 2.86, p = 0.02, n = 9 mice/group, unpaired t-test). (**C**) Evoked dopamine (Interaction effect: F(19,372) = 9.03, p < 0.0001; main effect of semaglutide treatment: F(1, 20) = 8.07, p = 0.01, main effect of drug application: F(2.82,55.15) = 38.42, p < 0.0001, n = 11 slices/5 mice/group, mixed-effects analysis) and (**D**) corresponding tau (Interaction effect: F(19,327) = 2.28, p = 0.002; main effect of semaglutide treatment: F(1, 18) = 7.80, p = 0.01; main effect of drug application: F(6.075, 104.6)= 14.76, p < 0.0001, n = 9-11 slices/5 mice/group, mixed-effects analysis) time courses with leptin (100 μM) application preceding the addition of cocaine (1 μM) *and* leptin (100 nM), normalized to the group average leptin response. (**E**) Evoked dopamine in response to leptin or leptin pretreatment followed by cocaine in mice previously treated with vehicle (Interaction effect: F(19,377) = 21.77, p < 0.0001; main effect of cocaine: F(1, 20) = 28.45, p <0.0001; main effect of drug application: F(2.78,55.10) = 41.72. p < 0.0001. n = 11 slices/3-5 mice/group. mixed-effects analysis) and (**F**) semaglutide (Interaction effect: F(19,333) = 3.67, p < 0.0001; main effect of cocaine: F(1, 23) = 0.45, p = 0.51; main effect of drug application: F(5.41,94.87) = 18.34, p < 0.0001, n - 11-14 slices/4-5 mice/group, mixed-effects analysis). (**G**) Analysis of dopamine reuptake (tau) following cocaine or leptin *and* cocaine application in fasted mice previously treated with vehicle (Interaction effect; F(19,329) = 1.64, p = 0.05; main effect of cocaine: F(1, 18) = 0.21 p = 0.65; main effect of drug application: F(4.61, 79.78) = 29.24, p < 0.0001, n = 9-11 slices/3-4 mice/group, mixed-effects analysis) or (**H**) semaglutide (Interaction effect: F(19,333) = 3.67, p < 0.0001; main effect of cocaine: F(1, 15) = 8.66. p = 0.01; main effect of drug application: F(5.41,94.87) = 18.34. p< 0.0001, n = 8-9 slices/3-5 mice/group, mixed-effects analysis). Data is presented as mean ± SEM; symbols are individual sample data. For all panels, *p < 0.05, ****p < 0.0001.

When we plotted the evoked dopamine following leptin vs. leptin and cocaine, the mice chronically treated with vehicle and fasted overnight showed a robust increase in cocaine-induced striatal dopamine compared to leptin-induced striatal dopamine alone (Figure 6E). However, mice chronically treated with semaglutide and fasted overnight showed no difference in evoked dopamine with leptin vs. leptin and cocaine bath applications (Figure 6F). This data demonstrates leptin’s restoration of semaglutide-induced regulation of cocaine in fasted mice, indicating the significance of leptin’s role in the mechanism. In control mice, there was no difference in dopamine reuptake following leptin or leptin and cocaine, while semaglutide-treated mice showed slower dopamine reuptake kinetics following leptin and cocaine vs. leptin alone (Figure 6G,H).

Together, these data suggest that leptin’s contribution to diminishing the cocaine response in semaglutide treatment is DAT-mediated.

## Discussion

### Chronic semaglutide inhibits the dopaminergic response to cocaine

Expanding on prior research demonstrating GLP-1 receptor agonists are effective in reducing cocaine use, our work provides novel insight into how chronic semaglutide reduces the rewarding effects of cocaine. Similar to recent findings on acute semaglutide administration [38], following chronic semaglutide, we demonstrate a reduction in the dopaminergic response to cocaine without inducing sedation effects. This study offers the first mechanistic insight into semaglutide’s inhibition of the dopaminergic and behavioral response to cocaine. DAT has been a proposed therapeutic target for CUD due to cocaine’s strong binding affinity, inducing reward signaling through dopamine reuptake antagonism [5–8]. Here, we show that chronic semaglutide caused a significant reduction in the dopamine released from cocaine through decreased inhibition of dopamine reuptake, aligned with previous research [61, 62]. Additional work has utilized DAT overexpression models to show that increased basal reuptake generated a decreased dopaminergic response to cocaine [63]. Some studies have suggested that increased striatal dopamine reuptake velocity may produce conditions more susceptible to cocaine inhibition, generating an elevated reward response and motivational state for cocaine [64, 65]. Despite heightened dopamine reuptake capability in baseline conditions, our semaglutide-treated mice displayed resistance to cocaine in FSCV and behavioral experiments. We propose that this phenomenon may be attributed to dependence on the contributions of striatal leptin’s regulation of DAT, as seen through the loss of semaglutide efficacy for cocaine regulation in a fasted state. Our findings reiterate the significance of DAT as a therapeutic target, propose a potential mechanism for GLP-1 receptor agonists’ regulation of addiction in leptin, and strengthen semaglutide’s case as a potential treatment option for chronic cocaine use.

### Semaglutide’s effects on cocaine are associated with increased striatal leptin sensitivity

In this study, we found that overnight fasting nullified the beneficial effects of semaglutide on cocaine despite long-term administration. By evaluating alterations in satiety hormones known to modulate the cocaine response, we determined leptin as a candidate for mediating semaglutide-induced inhibition of cocaine. We were able to replicate the finding of reduced fasting leptin concentrations in the periphery and decreased visceral fat in lean mice chronically treated with semaglutide [30].

In our FSCV and behavioral experiments, we also observed semaglutide-induced striatal leptin sensitivity for the regulation of dopamine release, reuptake, and locomotion. Leptin is known to generate a significant increase in dopamine release in the striatum through CIN activation, as well as an increase in dopamine reuptake by binding to its receptor present on dopaminergic neurons in the DMS [59]. Through application of DhβE, we isolated leptin signaling in dopaminergic neurons to observe the expected increase in dopamine reuptake upon leptin receptor activation. The reduction of tau in response to activation supports our hypothesis that leptin boosts reuptake of dopamine in semaglutide-treated mice. This effect may be driving semaglutide’s control over cocaine, as demonstrated by leptin rescuing the restriction in the cocaine response of fasted animals.

In our locomotion and FSCV experiments, mice given chronic semaglutide show particularly diminished responses to cocaine when pretreated with leptin after fasting overnight. As semaglutide has been shown to decrease visceral fat and fasting blood leptin concentrations in addition to increasing leptin sensitivity in the hypothalamus of obese models [46], we suspect that repeating these experiments in mice maintained on a high-fat diet would produce similar results. In some models, obesity causes increased sensitization to cocaine [21]. It is likely that semaglutide’s effects on dopamine and leptin during cocaine use would translate to a patient population with a range of body weights.

### Potential pathways connecting leptin to DAT for cocaine regulation

Previous research has demonstrated the significance of leptin for both DAT and cocaine regulation, as well as other markers downstream of leptin receptor activation. Specifically, extracellular signal-regulated kinase (ERK) phosphorylation produces DAT phosphorylation at Thr53, causing translocation to the synaptic cleft in which dopamine reuptake increases [66]. Importantly, ERK signaling facilitates leptin signal propagation within the striatum [67]. Investigating ERK as a target for increased DAT translocation to reduce the dopamine response to cocaine may lead to insight into how to directly influence striatal reward signaling in CUD patients. Additionally, the leptin signaling pathway leads to the inhibition of GSK3β, known to manipulate DAT trafficking [68] and cocaine sensitivity and behaviors [22, 69, 70]. Further exploration into the distinct molecular mechanisms generating leptin-induced cocaine insensitivity through semaglutide treatment could reveal the significant mediators of GLP-1’s reward regulation.

### Concern for weight loss in CUD patients on semaglutide

A common concern of the use of an anorexigenic drug for cocaine use disorder is the often observed reduction in fat mass and leptin in models and patients [71–74]. A study tracking the eating habits, weight, and leptin concentration of cocaine use disorder patients determined an increase in food intake with cocaine dependency despite weight loss and a trend in decreased leptin levels [75]. This confounding finding may be caused by elevated sympathetic nervous system activation from cocaine [76] that can reduce leptin secretion [77]. Self-administration of cocaine reduced glp-1 levels in rodents [78], and infusion of cocaine reduced GLP-1 levels in chronic users [79]. Peripheral GLP-1R activation was shown to reduce sympathetic nerve activity [80], adding to its potential for cocaine regulation and leptin signaling restoration in chronic users. It is also possible that GLP-1 treatment in cocaine use disorder may restore leptin production and signaling through its action on adipocytes [81]. Additionally, semaglutide may regulate substantial weight gain often seen in cessation of chronic cocaine use [82]. It would be of high importance to carefully monitor eating behaviors and leptin levels of CUD patients administered semaglutide.

### Experimental limitations and future directions

Importantly, semaglutide may also act on the hindbrain and arcuate nucleus of the hypothalamus GLP-1Rs to indirectly modulate striatal function [31, 83]. In support of this, previous research has demonstrated that GLP-1R expressing neurons in the nucleus tractus solitarius regulate cocaine seeking [78]. Our findings do not exclude contributions from hindbrain circuits, which may also be shaped by leptin-dependent or -independent mechanisms [84], a topic which requires future study.

Our findings are limited to male mice, but evidence supports testing this mechanism in females: female heterozygous DAT-knockout rats show increased locomotor sensitivity to cocaine [62], exogenous leptin reduces cocaine intake and accelerates extinction in female rats [20], and semaglutide lowers circulating leptin in female mice [85], consistent with our findings in males. We focused FSCV experiments on the dorsal striatum, where dopamine reuptake [39, 50] and its enhancement by leptin [59] are greater. Although this region contributes to cocaine-motivated behaviors and is altered in cocaine use disorder, future studies will examine semaglutide’s effects on cocaine and leptin sensitivity in the ventral striatum. Cocaine self-administration studies will assess relevance to chronic use, while striatal leptin receptor antagonism or knockdown will test whether leptin signaling is necessary for semaglutide’s effects.

## Acknowledgements

We gratefully acknowledge Dr. Chris Pierce for his contribution through expertise.

