## Supplementary Figure for "Semaglutide Increases Striatal Leptin Sensitivity, Impacting Dopaminergic Response to Cocaine"

### Supplementary Figures

#### Muldowney et al., Supplemental Figure 1

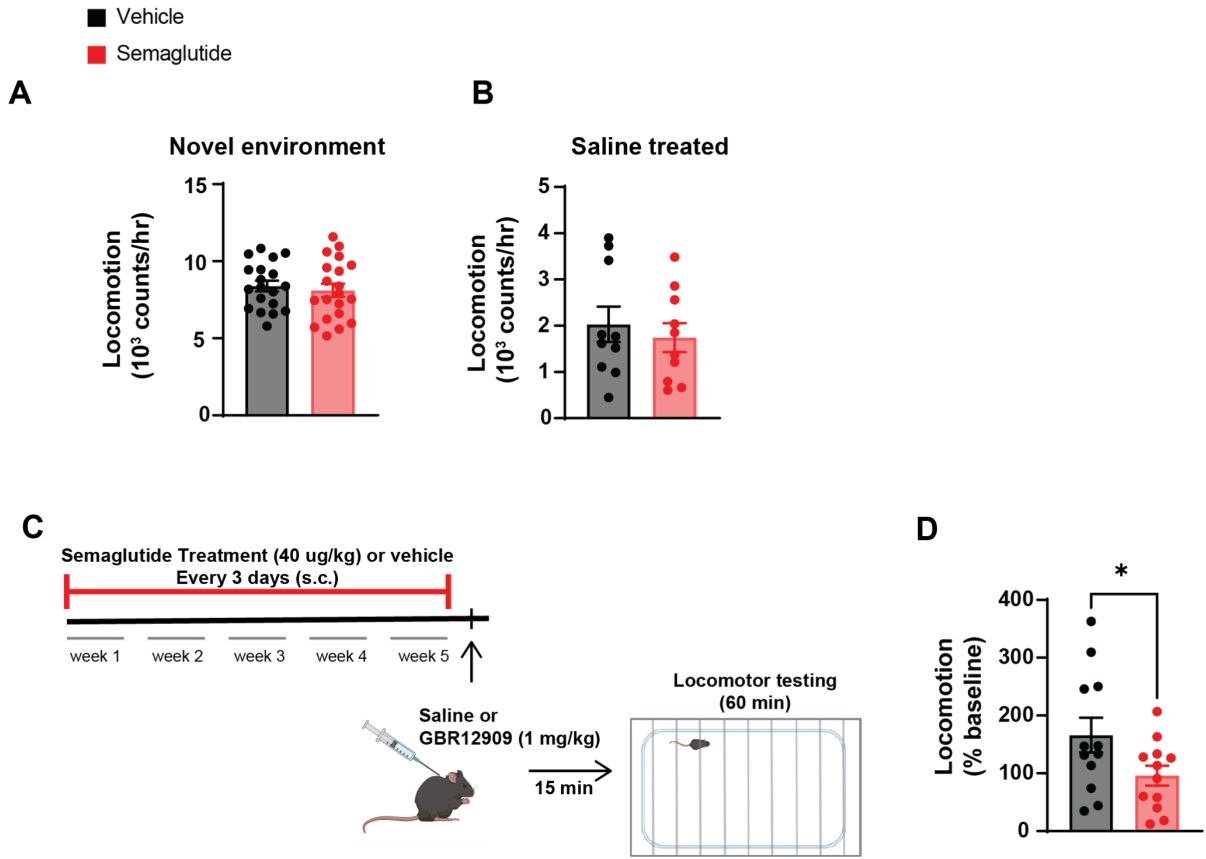

##### Supplementary Figure 1: Locomotion in novel environment, saline, and GBR trials.

(A) Locomotion per group during first exposure to locomotion boxes over a 60-minute period ($t(18) = 1.30$, $p = 0.21$, $n = 10$ mice/group, unpaired t-test). (B) Total locomotion during 60-minute trial following saline injection ($t(18) = 0.58$, $p = 0.57$, $n = 10$ mice/group, unpaired t-test). (C) Schematic demonstrating the injection protocol used in Supplementary Figure 1D. (D) Average locomotion of mice following injection of GBR 12909 (1 mg/kg) over a 60-minute trial, normalized to group-average saline control trial ($t(22) = 2.12$, $p = 0.04$, $n = 12$ mice/group, unpaired t-test). Data is presented as mean ± SEM; symbols are individual sample data. \*$p < 0.05$.

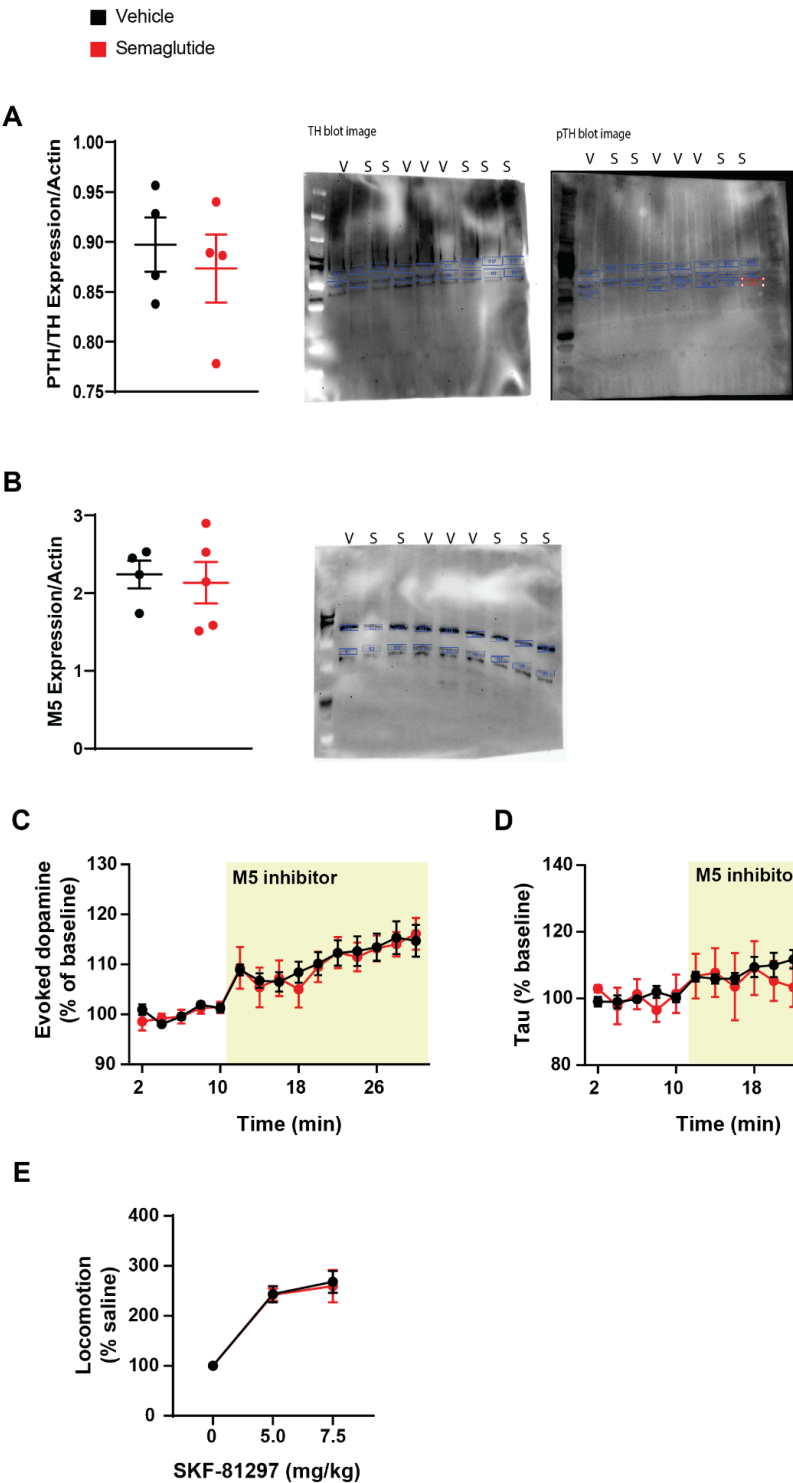

**Supplementary Figure 2: Analysis of additional dopaminergic signaling modulators.**

(A) Western blot analysis of the ratio of pTH to TH protein expression, normalized to actin (housekeeping gene) in dorsal striatal samples of fasted mice previously treated with either vehicle (V) or semaglutide (S) (t(6) = 0.55, p = 0.60, n = 4 mice/group, unpaired t-test). Images used for quantification of TH (left), phospho-TH (right), and actin. (B) Western blot analysis of M5 receptor expression in the dorsal striatum of vehicle and semaglutide-treated mice (t(7) = 0.31, p = 0.76, n = 4-5 mice/group, unpaired t-test). (right) Image used for quantification of M5 and actin. (C) Normalized evoked dopamine time course in response to the M5 inhibitor, ML375 (10 μM), application (Interaction effect: F(14,292) = 0.24, p = 0.99; main effect of semaglutide treatment: F(1,21) = 0.05, p = 0.83; main effect of drug application: F(3.66,76.26) = 19.65, p < 0.0001, n = 10-13 slices/4 mice/group, mixed-effects analysis). (D) Corresponding normalized tau time course in response to ML375 (10 μM) application (Interaction effect: F(14,273) = 0.57, p = 0.89; main effect of semaglutide treatment: F(1,20) = 0.002473, p = 0.96; main effect of drug application: F(5.94,115.8) = 4.41, p = 0.005, n = 9-13 slices/4 mice/group, mixed-effects). (E) Average locomotion per group in varying doses of a D1-like agonist SKF-81297 (Interaction effect: F(2,36) = 0.04, p = 0.97; main effect of semaglutide treatment: F(1,18) = 0.04154, p = 0.84; main effect of drug dosage: F(1.39,25.10) = 53.01, p < 0.0001, n = 10 mice/group, RM two-way ANOVA). Data is presented as mean ± SEM; symbols are individual sample data.

#### Muldowney et al., Supplemental Figure 3

**A**

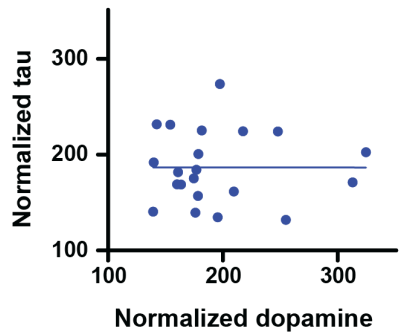

##### Supplementary Figure 3: Correlation between evoked dopamine and reuptake in fasted mice.

(A) Correlation of normalized dopamine and tau following cocaine application in brain slices from fasted mice ($R^2 = 0.06$, $p = 0.39$, $n = 15$ slices/6 mice, simple linear regression).

#### Muldowney et al., Supplemental Figure 4

**A**

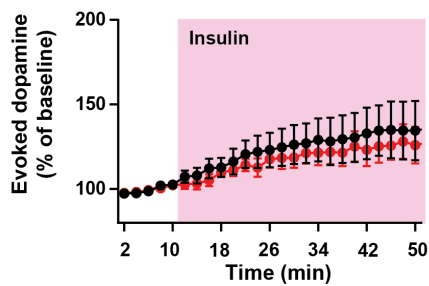

##### Supplementary Figure 4: Evoked dopamine following insulin application.

(A) Normalized evoked dopamine time course in response to insulin (100 nM) application (Interaction effect: $F(34,372) = 0.24$, $p > 0.99$; main effect of semaglutide treatment: $F(1, 11) = 0.23$, $p = 0.64$; main effect of drug application: $F(1.15, 12.52) = 5.48$, $p = 0.03$, $n = 5-8$ slices/3 mice/group). Data is presented as mean ± SEM.
